# Cerebellar Activity in PINK1 Knockout Rats during Volitional Gait

**DOI:** 10.1101/2023.02.28.530534

**Authors:** Valerie DeAngelo, Justin D Hilliard, Chia-Han Chiang, Jonathan Viventi, George C McConnell

## Abstract

Preclinical models of Parkinson’s disease are imperative to gain insight into the neural circuits that contribute to gait dysfunction in advanced stages of the disease. The PTEN-induced putative kinase 1 (P1) knockout (KO) early onset model of Parkinson’s disease may be a useful rodent model to study the effects of neurotransmitter degeneration caused by loss of P1 function on brain activity during volitional gait. The goal of this study was to measure changes in neural activity at the cerebellar vermis (CBLv) at 8 months of age. Gait deficits, except run speed, were not significantly different from age-matched wild-type (WT) controls as previously reported. P1KO (*n*=4) and WT (*n*=4) rats were implanted with a micro-electrocorticographic array placed over CBLv lobules VI (a, b, and c) and VII. Local field potential recordings were obtained during volitional gait across a runway. Power spectral analysis and coherence analysis were used to quantify network oscillatory activity in frequency bands of interest. CBLv power was hypoactive in the beta (VIb, VIc, and VII) and alpha (VII) bands at CBLv lobules VIb, VIc, and VII in P1KO rats compared to WT controls during gait (p<0.05). These results suggest that gait improvement in P1KO rats at 8 months may be a compensatory mechanism attributed to movement corrections caused by decreased inhibition of the alpha band of CBLv lobule VII and beta band of lobules VIb, VIc, and VII. The P1KO model may be a valuable tool for understanding the circuit mechanisms underlying gait dysfunction in early-onset Parkinson’s disease patients with functional loss of P1. Future studies investigating the CBLv as a potential biomarker and therapeutic target for the treatment of gait dysfunction in Parkinson’s disease are warranted.

## Introduction

Animal models of Parkinson’s disease are important for understanding the pathophysiology during disease progression and for the development of promising treatments. An ideal model of Parkinson’s disease should be age dependent and progressive.^1^ The PTEN-induced putative kinase 1 (P1) knockout (KO) in which the P1 gene is deleted, is a commercially available rat model for early onset Parkinson’s disease.^2–3^ It is unclear if this model may be beneficial for understanding the circuit mechanisms involved in Parkinsonian gait dysfunction.

A pathological hallmark of Parkinson’s disease is dopamine (DA) degeneration in the substantia nigra pars compacta (SNc) and cholinergic degeneration in the pedunculopontine nucleus (PPN).^4^ An early study of this model measured 50% DA depletion in the SNc when compared to healthy wild type (WT) rats and locomotor deficits beginning as early as 4 months and continuing up to 8 months of age.^5^ Succeeding studies, however, presented mixed results. Reports of DAergic degeneration ranged from no loss^6–7^ to significant loss^8–9^, gait deficits showed improvement at 8 months of age^5,10^, and no significant loss of acetylcholine was found in the PPN.^10^

Electrophysiological measurements of brain regions associated with abnormal gait may provide insight into the mechanisms of gait dysfunction in patients associated with loss of P1 function. Motor behaviors are associated with coordinated neural networks that are distributed across functionally connected brain regions. Local field potential (LFP) recordings from various brain regions allow insight into the circuit dysfunction contributing to specific symptoms such as abnormal gait.^11^

Voluntary movement control and execution are regulated by cerebellar influence over different interconnected cortical areas.^12^ Multimodal inputs to the fastigial nucleus (FN) coordinate postural responses by sending bodily information during walking to posture and gait related areas in the brainstem and cerebral cortex.^13^ The cerebellar vermis (CBLv) is mediated by FN output neurons and is hyperactive in Parkinson’s disease^2–3^, making it a possible contributor to deficits exhibited during Parkinsonian gait.

We hypothesized that pathological cerebellar activity contributes to gait dysfunction in the P1KO model of Parkinson’s disease. Using a micro-electrocorticographic (μECoG) array, LFP activity was recorded from the CBLv during gait in P1KO rats at 8 months of age. Power spectral analysis and coherence analysis were used to analyze μECoG recordings and quantify functional connectivity between lobules compared to age-matched WT controls.

## Materials and Methods

### Animals and housing

Adult male Long-Evans hooded P1KO rats (*n* = 4) and age-matched Long-Evans hooded WT rats (*n* = 4) derived from the breeding of the P1KO rats were purchased from Horizon Discovery SAGE labs. Rats were singly housed (post-implantation surgery) in standard cages with free access to food and water. Study protocols were reviewed and approved by the Stevens Institute of Technology Institutional Animal Care and Use Committee. These rats were used in a previous study.^10^

### Gait analysis

Prior to implantation gait assessment was performed on P1KO and age matched WT controls using the Cleversys runway system (CSI-G-RWY; CleverSys Inc., Reston, VA). As the rat voluntarily moves across the runway, comprised of a long side-lit glass plate, each footprint is lit up and recorded by a camera mounted under the glass. Measurements obtained were swing time (time in which paw is in the air), stance time (time in which all paws are detected on the glass), stride time (stance time plus swing time), base of support (average width between hind paws), stride length (distance the paw traversed from start of previous stance to beginning of next stance) and run speed (instantaneous speed over a running distance).^10^

### Implantation of μECoG array

When rats reached 8 months of age, sterile stereotactic surgery was conducted under 4% sevoflurane anesthesia using coordinates from a rat brain atlas by Paxinos and Watson.^14^ A 5mm x 6mm craniotomy was created over CBLv (AP −9mm to 14mm, ML ±3mm) (Fig. 1A). Each rat was implanted with a μECoG array (Fig. 1B) arranged in a 8×8 grid with 203μm diameter spaced 406μm apart over lobules VIa (AP −11.4mm to −12mm, ML ±1.3mm), VIb (AP −12.2 mm to −12.8mm, ML ±1.3mm), VIc (AP −13 mm to −13.2mm, ML ±1.3mm), and VII (AP −13.4mm to −13.6mm, ML ±1.3mm) (Fig. 1C). Following electrode implantation, sterile Covidien Vaseline petroleum jelly was placed over the craniotomy to create a barrier between the head cap and the brain. This prevented potential damage to the vermis through direct contact with the dental cement used to create the headcap.

**Figure 1:**
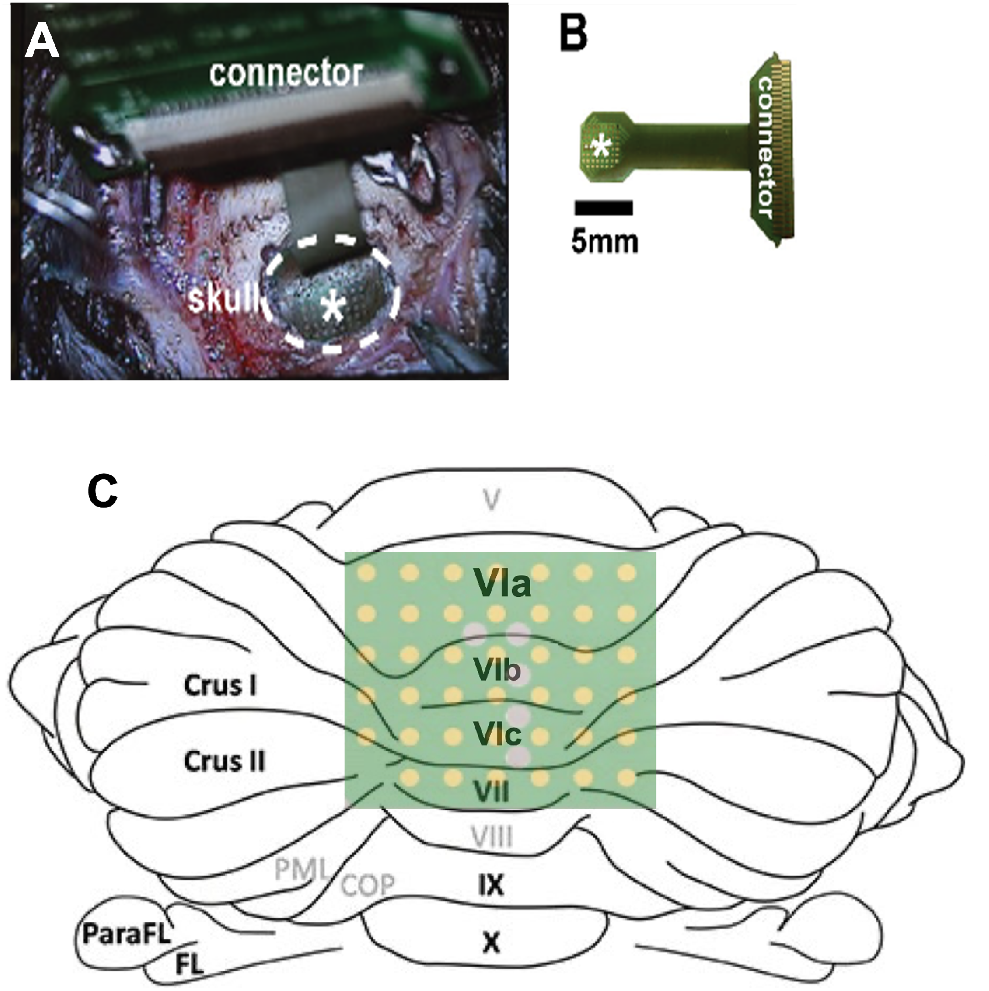
μECoG implantation and placement. (A) Microscopic image of μECoG placed over the vermal craniotomy made using a 5mm trephine. (B) Dimensions of the μECoG array and connector. * indicates electrode contacts. (C) Approximate placement of electrodes over cerebellar vermis lobules VIa, VIb, VIc, and VII.

### Neural data acquisition

One week post-implantation, LFP recordings were taken during volitional gait on the runway using the Open Ephys (open-source electrophysiology) system (Fig. 2A-B). The runway was modified to synchronize LFP recordings with gait. A gap was created from the start box to the exit box to allow the serial peripheral interface (SPI) cable, connecting the headstage to Open Ephys, to move freely with the rat. Two beam break sensors were used in tandem with an Arduino to control the LFP record timing. As the rat left its start box and entered the field of view, the first sensor was broken and a TTL pulse was sent to Open Ephys to start recording. LFPs were continuously recorded as the rat traversed the runway. After the rat exited the runway and entered the exit box, a second sensor sent a pulse to stop the recordings. Data from three consecutive runs were averaged for each rat. Gait was assessed prior to implantation in a previous study.^10^

**Figure 2:**
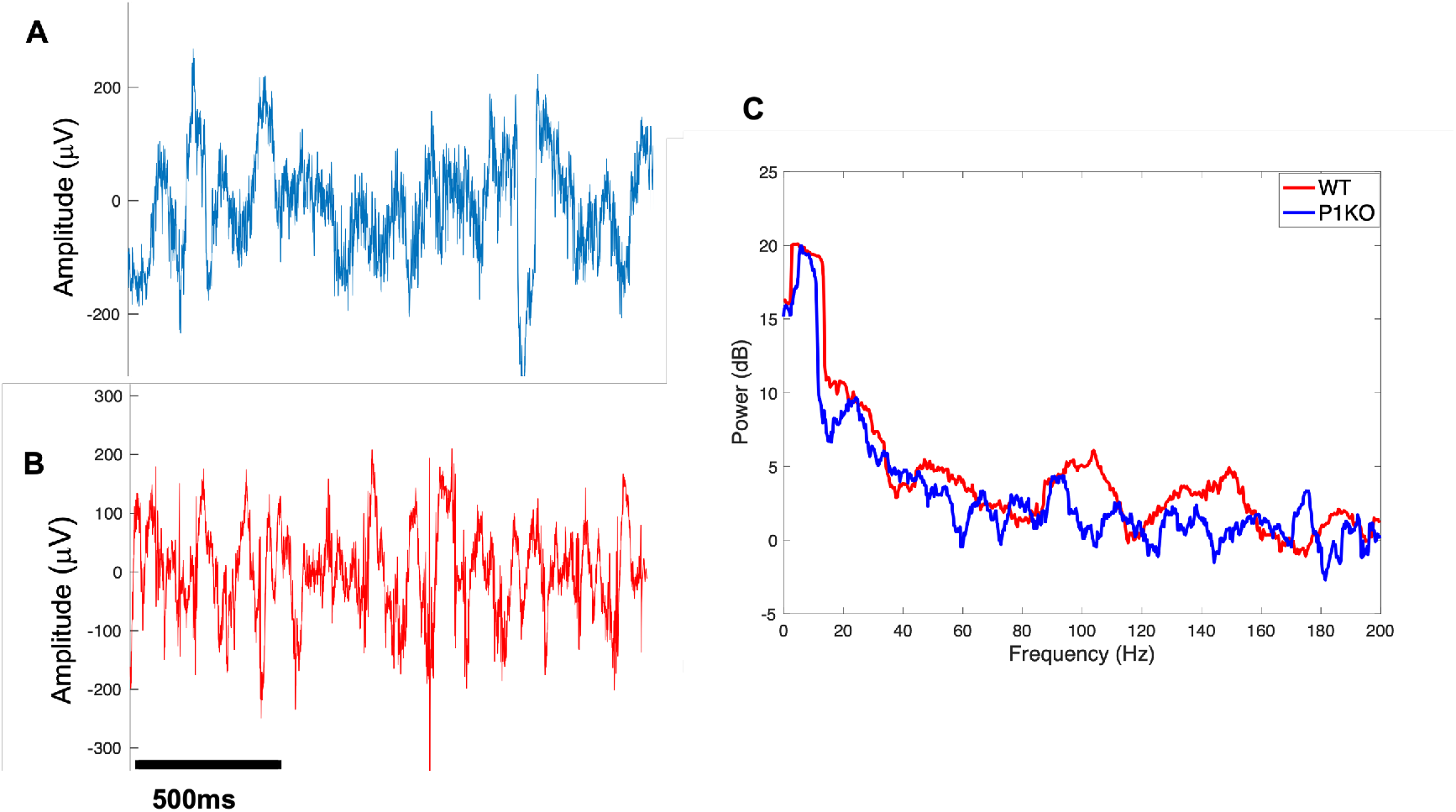
Sample μECoG recordings. (A) Sample LFP recorded from cerebellar vermis lobule VIa of P1KO rat and (B) WT rat. (C) Sample power spectra of LFPs recorded from P1KO and WT rats during volitional gait.

### Neural data analysis

Continuous LFP recordings were exported and analyzed using MATLAB (Mathworks, Natick, MA, United States). Spectral analysis was used to quantify changes in CBLv oscillatory activity Multitaper methods of spectral estimation were used to quantify changes in CBLv oscillatory activity between the P1KO and WT genotype in each lobule (Chronux version 2.00) (Fig. 2C). The LFP spectrum was estimated on a 2 second window with 10 Hz resolution using 19 Slepian data tapers.^23^ The mean power in dB was measured within the following frequency bands: delta (1-4Hz), theta (4-8 Hz), alpha (8-13 Hz), low beta (13-20 Hz), high beta (21-30 Hz), low gamma (30-55 Hz), high gamma (65-80 Hz) and fast frequency (80-200 Hz). Frequencies between 55 Hz and 65 Hz were omitted to reduce effects of 60 Hz noise.^16^ Coherence analysis was performed at the same frequencies to determine functional connectivity between different lobules of the CBLv. The mean coherence between electrodes in all possible pairs of vermis lobules was measured and compared at each frequency band. Coherence was estimated on a 2 second window with 5 Hz resolution using 19 Slepian data tapers.^23^

### Immunohistochemistry

Immediately following LFP recordings, rats were deeply anesthetized and transcardially perfused with PBS followed by 10% formalin. The brain was removed and postfixed overnight (4°C) in formalin then placed in a 30% sucrose (4°C) solution until it sank. A green tissue marking dye was applied to the left-posterior hemisphere to demarcate the orientation of the brain sections. The brains were cryoprotected with Tissue-Tek optimal cutting temperature (O.C.T.) compound and 40μm serial coronal sections were cut using a cryostat (CryoStar NX50) equally spaced through the SNc, striatum, and pedunculopontine nucleus (PPN). Immunohistochemistry was performed in the SNc and striatum with anti-tyrosine hydroxylase (TH) antibody to measure DAergic neuron loss and on the PPN with anti-choline acetyltransferase (ChAT) antibody to measure cholinergic neuron loss. Details of staining and cell counting were performed as reported by DeAngelo et al.^10^

### Statistical analysis

Statistical analysis was performed using IBM SPSS Statistics for Mac (IBM Corp., Armonk, N.Y., USA). A one-way ANOVA was used to determine significance between genotype and average power in each lobule and differences in average power between lobules. An independent Student’s *t*-test was used to detect significant differences in lobule pair (interlobular) coherence between groups. Interlobular coherence within each genotype was analyzed using a oneway ANOVA and Tukey’s post hoc test to detect significant differences between frequency bands. If homogeneity of variances was violated a Welch ANOVA with Games-Howell post hoc test was used. Results were considered statistically significant at *p* ≤ 0.05. All bars show mean +/-SEM.

### Data availability

Raw data were generated at Stevens Institute of Technology. Derived data supporting the findings of this study are available from the corresponding author on request.

## Results

### Gait analysis and cell counting

Results were previously reported.^10^ Briefly, at 8 months of age, gait measurements in P1KO rats more closely resembled those found in WT, while WT remained unchanged over time. P1KO mean stance time, stride time, swing time, stride length, and run speed decreased and base of support stayed the same. Aside from stride length, these changes were not significant compared to the same measures at 5 months. Run speed was the only measure that was significantly lower than WT controls (*p*<0.05) at 8 months. Cell counting resulted in a 26.7% loss of TH+ cells in the SNc of P1KO compared to WT. No significant TH+ terminal loss or ChAT+ cells were reported in the striatum and the PPN, respectively.

### Neural data analysis: Power

This study was unblinded. LFP mean logarithmic power was measured in CBLv lobules VI (a, b, and c) and VII in the delta, theta, alpha, low beta, high beta, low gamma, high gamma, and fast frequency bands in P1KO rats and WT controls. Vermis lobules of P1KO rats were hypoactive when compared to WT rats in all frequency bands in all lobules.

Significant differences were found in the alpha, low beta, and high beta bands. Hypoactivity in the P1KO model was measured as significant in the alpha band of lobule VII (*p*=0.048), low beta band of lobules VIb (*p*=0.005), VIc (*p*=0.03), and VII (*p*=0.027) and high beta band of VIb (*p*=0.021), VIc (*p*=0.027), and VII (*p*=0.031) when compared to WT controls (Fig. 3).

**Figure 3:**
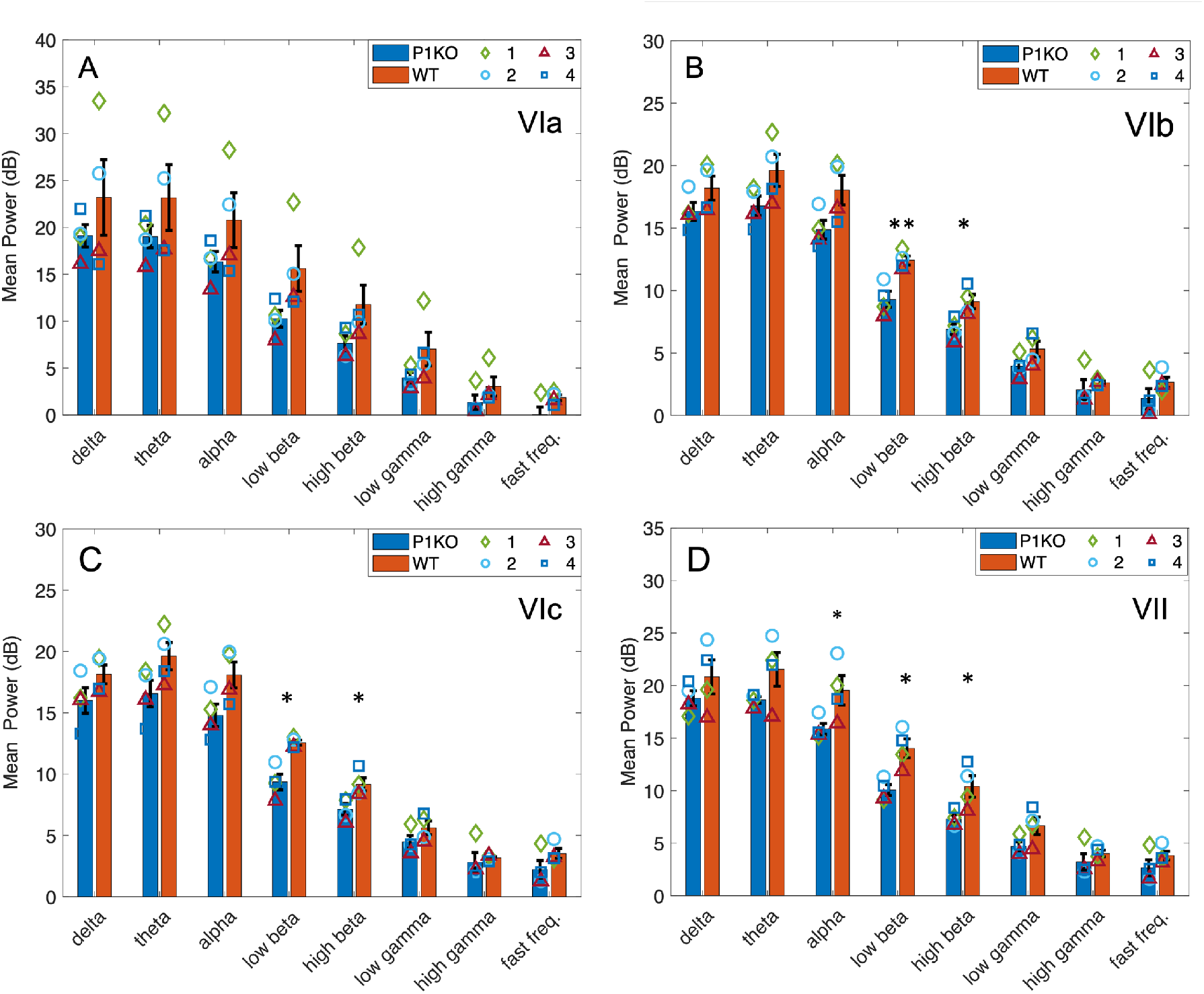
Spectral power analysis. (A) No significant differences were measured in power between genotypes in lobule VIa. P1KO rats had significantly lower LFP activity in the low beta and high beta bands in (B) lobules VIb, (C) VIc, and (D) VII. P1KO was hypoactive compared to WT across all bands in all lobules. Symbols represent the mean power in each band in P1KO (*n* = 4) and WT rats (*n* = 4). * indicates *p*<0.05; ** indicates *p*<0.01.

Power between lobules in each frequency band, measured using a one-way ANOVA, showed similar activity in frequency bands below 65Hz in all lobules in both P1KO and WT (*p*>0.1). WT rats exhibited significant differences between lobular activity in the high gamma (*p*=0.022) and fast frequency (*p*=0.037) bands. Post hoc analysis revealed greater high gamma activity in lobules VIc (*p*=0.045, Fig. 3C) and VII (*p*=0.05, Fig. 3D) compared to lobule VIb (Fig. 3B) and fast frequency in lobules VIc (*p*=0.041) and VII (*p*=0.016) compared to VIa (Fig. 3A).

### Neural data analysis: Coherence

Mean coherence was measured between each pair of recording sites in the delta, theta, alpha, low beta, high beta, low gamma, high gamma, and fast frequency bands in P1KO rats and WT controls (Fig. 4). Pairings were between lobules: VIa and VIb; VIa and VIc; VIa and VII; VIb and VIc; VIb and VII; VIc and VII. Varying degrees of coherence were observed across pairs with the greatest coherence in neighboring lobules, which decreased with interelectrode distance.

**Figure 4:**
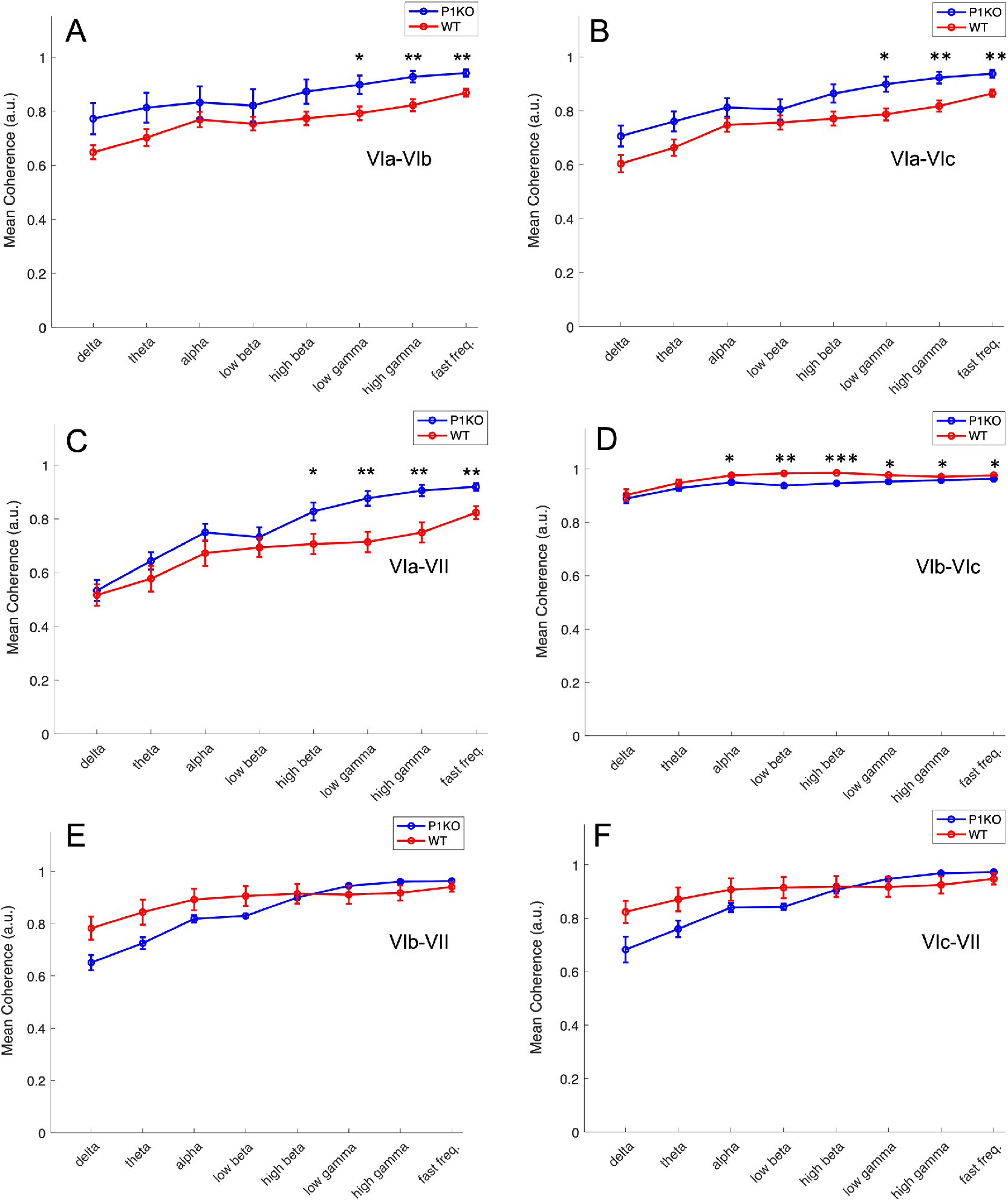
Coherence analysis. Interlobular coherence was calculated in each frequency band. Coherence increased in P1KO (*n*=4) compared to WT rats (*n*=4) in lobular pair: (A) VIa-VIb and (B) VIa-VIc in low gamma, high gamma, and fast frequency bands. (C) P1KO had greater coherence than WT between lobules VIa and VII in the high beta, low gamma, high gamma, and fast frequency bands. (D) WT rats had significantly greater coherence in lobules VIb and VIc in the alpha, low beta, high beta, low gamma, and fast frequency bands. No significant differences were found in pairs (E) VIb-VII and (F) VIc-VII. * indicates *p*<0.05; ** indicates *p*<0.01; *** indicates *p*<0.001.

Comparison of mean coherence between P1KO and WT showed similar activity (Fig. 4) at lower frequencies (i.e., delta, theta, alpha, low beta, and high beta) in most pairings except for: 1) VIb-VIc (Fig. 4D), where coherence significantly decreased in P1KO rats in the alpha (*p*=0.023), low beta (*p*=0.001) and high beta (*p*<0.001) bands and 2) VIa-VII (Fig. 4C), where P1KO coherence significantly increased in the high beta band (*p*=0.028). Coherence decreased in WT rats vs. P1KO rats between pairs VIa–VIb, VIa-VIc, and VIa-VII (Fig. 4A-D) for the low gamma (*p*=0.011; *p*=0.032; *p*=0.006), high gamma (*p*=0.006; *p*=0.007; *p*=0.005), and fast frequency (*p*=0.004; *p*=0.005; *p*=0.006) bands. Coherence increased in WT rats vs. P1KO rats between lobules VIb and VIc at low gamma (*p*=0.010), high gamma (*p*=0.042), and fast frequency (*p*=0.014) bands.

Interlobular coherence was directly related to frequency in both genotypes. P1KO coherence significantly differed between frequency bands in all pairings (*p*<0.005) except for VIa-VIb (*p*=0.153). Post hoc analysis revealed increased coherence between electrodes in the low gamma, high gamma, and fast frequency bands compared to the delta, theta, alpha, low beta and high beta bands for all lobule pairings except for VIb-VIc (Fig. 4C). Coherence in the low gamma band was greater than delta (*p*<0.05), theta (*p*<0.05), alpha (*p*<0.05) and low beta bands (*p*<0.05) between electrodes in VIb-VII and VIc-VII. Pairing VIa-VII significance excluded alpha (*p*=0.093) and between VIa-VIc low gamma was only greater than the delta band (*p*=0.003). Both high gamma and fast frequency bands were significantly greater than delta, theta, alpha and low beta (*p*<0.05) in all pairings except for VIa-VIc which exhibited significance over the delta (*p*<0.001) and theta (*p*<0.02). Coherence between VIc-VII in high gamma and fast frequency were also greater than coherence in the high beta band (*p*=0.05 and *p*=0.037, respectively).

Interlobular coherence in WT rats increased with frequency in all pairings except VIb-VIc with significant differences measured between VIa-VIb (*p*<0.001), VIa-VIc (*p*<0.001), and VIa-VII (*p*<0.001). Coherence was lowest in the delta band when compared to all other frequency bands with varying significance (Supplementary Table 1). Pairs VIa-VIb and VIa-VIc exhibited higher gamma and fast frequency coherence than delta and theta (*p*<0.05). Fast frequency coherence was also greater than low beta (*p*=0.046) in VIa-VIb, alpha (*p*=0.040) in VIa-VIc, and theta (*p*<0.001) in VIa-VII. In contrast to this pattern, coherence between lobules VIb and VIc was significantly increased in the beta band when compared to higher frequency bands. Low beta band coherence was greater than high gamma coherence (*p*=0.03) while high beta band coherence was greater than low gamma (*p*=0.035), high gamma (*p*=0.004), and fast frequency (*p*=0.033).

Comparison of interlobular coherence between regions was used to determine the degree of functional connectivity between lobules. Significant differences were detected in the P1KO model between all pairings in delta (*p*<0.001), theta (*p*<0.001), alpha (*p*<0.001), low beta (*p*<0.001), high gamma (*p*=0.046) high beta (*p*=0.006), and fast frequency (*p*=0.01). Significant differences were also detected in the WT model between all frequency bands (*p*<0.001). Coherence between lobular pair VIb-VIc was greater than 0.9 in all frequency bands in WT rats and all frequency bands except delta in P1KO rats.

Post hoc analysis in P1KO found significant differences between lobule pair VIb-VIc (*p*=0.019)and VIa-VIc (*p*=0.019), VIa-VII (*p*<0.001), VIb-VII (*p*=0.03), and VIc-VII (*p*=0.018) in the theta band, VIa-VII (*p*=0.008; *p*=0.014), VIb-VII (*p*<0.001; *p*<0.001), and VIc-VII (*p*=0.005; *p*=0.001) in the alpha and low beta bands, and VIb-VII (*p*=0.015) in the high beta band. No pairwise significant differences were measured between VIb-VIc coherence and other pairs in the high gamma and fast frequency bands. Significant differences were also found between VIa-VIb and VIa-VII (*p*=0.017) in the theta band and VIc-VII and VIa-VII (*p*=0.013) in the fast frequency band.

These differences were not as widespread in the WT rats. Post hoc analysis found coherence between VIb-VIc was greater than VIa-VIb (*p*≤0.005), VIa-Vc (*p*≤0.005), and VIa-VII (*p*<0.01) in all frequency bands. VIb-VII coherence was greater than VIa-VII in all frequency bands except alpha (*p*<0.05) and greater than VIa-VIc in the theta band (*p*=0.023). Increased coherence was observed between electrodes in VIc and VII in comparison to pair VIa-VII in all frequency bands (*p*<0.05), pair VIa-VIc in the delta (*p*=0.002) and theta (*p*=0.007) bands, and pair VIa-VIb in the theta band (*p*=0.040).

## Discussion

The exact mechanism of gait dysfunction in Parkinson’s disease is unknown but is associated with neurotransmitter abnormalities which cause disruptions in connections between the BG, PPN, and cerebellum^4^. DAergic cell loss measured in the SNc of P1KO rats may cause hyperactive oscillations in the CBLv associated with abnormal gait. A μECoG array was implanted over CBLv lobules VIa, VIb, VIc, and VII to measure LFP activity in P1KO and aged-matched WT controls.

P1KO rat vermis activity was hypoactive across all lobules and frequency bands. Power analysis revealed significant differences in the alpha, low beta, and high beta bands of lobules VIb, VIc, and VII. CBLv lobule VI receives inputs to and from both the premotor (PM) and primary motor cortices [M1].^2,17^ A disynaptic pathway exists connecting lobule VII to the subthalamic nucleus (STN) of the basal ganglia (BG) via the PPN making it susceptible to abnormal activity caused by DA loss in the SNc.^18–19^ Together lobules VI and VII are involved in cognition, emotion, and motor planning.^17,20^

Previous studies report that decreased CBLv inhibitory activity results in hypoactivation of Purkinje cells (output cells of the CBLv) when movement corrections are required to respond to environmental stimuli.^2,21–22^ Decreased CBLv inhibition also occurs in response to error signals implicating its contribution in motor adaptation.^23^ We speculate that gait improvement in P1KO rats from 5 months to 8 months of age may be due to movement corrections during gait facilitated by decreased inhibition in lobules VIb, VIc, and VII.

Hypoactivation of lobule VI (b and c) was measured in the beta frequency band. Excessive beta activity (12-30Hz) is a prominent feature in STN recordings in patients with Parkinson’s disease.^24–29^ Since the BG and cerebellum are connected in a functional loop, decreased oscillatory activity in the vermis could result from increased oscillatory activity in the STN.^3,19^ These results indicate that the mechanism by which gait is improved in P1KO rats is by excessive inhibition of the beta band, which may be a compensatory response to increased beta activity in the STN caused by DA depletion in the SNc. Further research measuring LFP activity in the BG and cerebellum of P1KO rats presenting significant gait impairments are needed to test this hypothesis.

Abnormal activity in lobule VII was also measured in the alpha band. Alpha oscillations are associated with attention and processing.^30^ Suppression of this band is thought to reduce the ability for smooth execution of motor commands.^31^ A study by Thevathasan et al. found that alpha band activity in the PPN had a strong positive correlation to gait speed.^31^ All deficits in gait function measured in P1KO rats improved from 5 months to 8 months of age except for run speed which remained significantly decreased compared to WT controls. Thus, the sustained reduction in gait speed in P1KO rats at both time points may be due to reduced alpha band activity of the CBLv. Lobule VIa in P1KO rats did not exhibit significant abnormal alpha band activity compared to WT. A recent study by Fujita et al. found that the dominant projection from the FN to lobule VIa was different from the FN subdivision that projects to the posterior region of lobule VI and to lobule VII^32^, which may explain the association between changes in alpha band activity with lobule VII but not VI.

To gain further understanding into the lobular mechanisms contributing to gait in the P1KO model, we used coherence to measure functional connectivity between lobules. Coherence analysis measured high functional connectivity between all lobules with the greatest connectivity between lobules VIb and VIc in both P1KO rats and WT controls. The functional connectivity between lobules VIb and VIc was significantly decreased in P1KO compared to WT controls. In rat, CBLv lobules VIb and VIc receive inputs from the same brainstem nuclei.^33^ A decrease in functional connectivity within these lobules may contribute to gait deficits present in this model. Coherence between lobules VIa and VIb was significantly lower than coherence between VIb and VIc in WT rats, supporting the finding that lobules VIb and VIc are more functionally connected than lobules VIa and VIb.^32^ Differences between these pairs in P1KO rats, however, was not significant. This change in coherence may be indicative of the disparity in hypoactivation of this lobule among P1KO rats.

Interlobular coherence was directly related to location and frequency in both genotypes. Coherence was reduced as pairings between lobules spaced further apart (e.g., coherence between lobules VIa-VIb was greater than between lobules VIa-VII). Coherence increased with increasing frequency. Purkinje cell activity is synchronized with high frequency LFP oscillations between 100 Hz and 200 Hz.^34–35^ Increased coherence at higher frequencies may be a response to this activity since Purkinje cells are the sole output neurons to the cerebellar cortex.

In conclusion, the results of this study shed light on the circuit mechanisms underlying gait impairment in the P1KO rat model of Parkinson’s disease. CBLv activity in lobules VI (b and c) and VII provide direct evidence for improvements in gait. Specifically, these results suggest that hypoactivity of the beta band in lobules VIb, VIc, and VII is a result of compensatory mechanisms by the CBLv to counteract abnormal BG activity and restore normal gait. This compensatory mechanism is broken down in the alpha band of lobule VII which may explain why run speed is reduced in P1KO rats compared to WT. A decrease in functional connectivity between lobules VIb-VIc may also contribute to these findings.

The P1KO rat model may be a valuable model to investigate the neural basis for gait dysfunction and novel treatments in early-onset Parkinson’s disease. We hypothesize that the CBLv compensates for gait dysfunction in early stages of Parkinson’s disease by decreased inhibition of Purkinje cells. Worsening of gait in the advanced stages of Parkinson’s disease may be explained by the inability of the CBLv to compensate for BG pathological oscillatory activity resulting in hyperactivity of the CBLv.^2–3,36–38^ Future clinical studies in Parkinson’s patients presenting gait dysfunction are warranted to investigate the utility of the CBLv as a biomarker for gait dysfunction and/or potential therapeutic target to treat

## Abbreviations

CBLv: cerebellar vermis
BG: basal ganglia
PPN: pedunculopontine nucleus
SNc: substantia nigra pars compacta
STN: subthalamic nucleus
DA: dopamine
DAergic: dopaminergic
FN: fastigial nucleus
LFP: local field potential
μECoG: micro-electrocorticographic
P1KO: PTEN-induced putative kinase 1 knock-out
TH: anti-tyrosine hydroxylase
CHAT: anti-choline acetyltransferase

