## Supplementary Table 1 for "Cerebellar Activity in PINK1 Knockout Rats during Volitional Gait"

### Supplementary Material

**Supplementary Table 1: Interlobular coherence.** Coherence was lowest in the delta band when compared to other frequency bands in WT rats. P-values of these interactions range from <0.001 to 0.03.

| Lobular Pair | Pairwise Comparisons (a>b) | | p-value |
|---|---|---|---|
| | **a** | **b** | |
| **VIa-VIb** | theta | delta | 0.030 |
| | high beta | | 0.022 |
| | low gamma | | 0.005 |
| | high gamma | | <0.001 |
| | fast freq. | | <0.001 |
| **VIa-VIc** | alpha | delta | 0.005 |
| | low beta | | 0.003 |
| | high beta | | <0.001 |
| | low gamma | | <0.001 |
| | high gamma | | <0.001 |
| | fast freq. | | <0.001 |
| **VIa-VII** | high beta | delta | 0.029 |
| | low gamma | | 0.020 |
| | high gamma | | 0.003 |
| | fast freq. | | <0.001 |
